# Cholesterol metabolism modulation facilitates CAR-T induced killing of ovarian cancer

**DOI:** 10.1101/2025.08.20.671345

**Authors:** Chinmayee V. Prabhu Dessai, Zhuoying Huang, Haonan Lin, Guangrui Ding, Hongjian He, Menna Siddiqui, Meng Zhang, Wilson Wong, Ji-Xin Cheng

## Abstract

Ovarian cancer is one of the most lethal gynecological cancers worldwide and has one of the highest recurrence rates. Recently developed Chimeric Antigen Receptor T (CAR-T) cell therapy has shown potent clinical efficacy against hematological malignancies. However, solid tumors, including ovarian cancer, possess several mechanisms that hinder T cell activity, and metabolic alteration of cancer cells has been shown to contribute to resistance to immune cell attack against solid tumors. Here, we explored the metabolic response of ovarian cancer cells to CAR-T cell attack using label-free high-content hyperspectral stimulated Raman scattering (h^2^SRS) imaging. Utilizing visible h^2^SRS imaging with much improved spatial resolution, we found an altered cholesterol metabolism, featured by increased storage of cholesteryl ester in lipid droplets and free cholesterol, in ovarian cancer cells that survived the CAR-T treatment. Administration of Avasimibe, an inhibitor of cholesteryl esterification, further enhanced CAR-T cytotoxicity. Our study shows the promise of implementing metabolic modulation to facilitate CAR-T cell treatment of solid tumors.

**Significance Statement:** Ovarian cancer is a major global health challenge, mainly due to relapse driven by chemotherapy resistance development. Meanwhile, immunotherapies such as CAR-T have revolutionized treatment for blood cancers, but their application in solid tumors like ovarian cancer is hindered by limited efficacy. This study uses high-content stimulated Raman scattering (SRS) imaging to reveal metabolic adaptations that help ovarian cancer cells survive CAR-T–mediated cytotoxicity. Guided by these insights, we apply metabolic interventions that enhance CAR-T cytotoxicity efficacy. This approach reveals therapeutic vulnerabilities in ovarian cancer and demonstrates an investigative strategy applicable for overcoming resistance in solid tumors.

## Introduction

Ovarian cancer poses a major clinical challenge and remains the leading cause of death among the gynecological malignancies in the world. Despite extensive research and treatment advances, it remains a persistent problem because of the high recurrence rates (1). The current standard of care for ovarian cancer includes surgical removal followed by platinum-based chemotherapy (2). Even though many patients show an initial positive response, the majority of them show a relapse within two years, most commonly due to platinum resistance development (2, 3). Due to this short-lived success of the existing standard therapy, there is an urgent need for more effective clinical strategies. A particularly exciting form of immunotherapy is the Chimeric Antigen Receptor (CAR) T cell therapy, which has demonstrated potent efficacy in the clinics against hematological malignancies (2). CAR-T cell therapy involves genetically engineering the patient’s T cells to express the CAR that binds to a specific antigen expressed on the target cancer cells. While CAR-T therapy has shown success in hematological malignancies, extending CAR-T therapy to solid tumors, such as ovarian cancer, has been challenging (4, 5). One of the major obstacles is the immunosuppressive tumor microenvironment, which limits the activity of CAR-T cells against solid tumors (5, 6). Hence, there is a need to improve CAR-T cell-mediated killing of cancer cells.

Cell metabolism plays an important role in CAR-T cell cytotoxicity and is pivotal in understanding immunotherapy (7–10). Similarly, modulation of cancer cell metabolism has also been linked to cancer survival under drug resistance and stress resistance (3, 11). The inhibition of several metabolic pathways has been shown to reduce cancer cell aggression (3, 12, 13). The metabolic dynamics of CAR-T cells attacking cancers will be an important piece of information for understanding the interaction between the engineered immune cells and tumors, and for discovering metabolic targets to improve the CAR-T cell-mediated killing efficacy. Current gold standard techniques to study cell metabolism include mass spectrometry and fluorescence imaging (14). Although highly sensitive, mass spectrometry-based methods do not elucidate metabolic dynamics or capture cellular heterogeneity due to their bulk analysis and destructive nature. Imaging mass spectrometry can provide spatial metabolic information but often requires sample preparation and is affected by limited spatial resolution. In contrast, fluorescence imaging is high-resolution but requires prior knowledge of the metabolic pathways to target, capping its applications in exploratory studies. Additionally, fluorescent probes are usually bulky and may interfere with the metabolic processes being investigated, potentially introducing artifacts in the data (15, 16). Label-free, non-destructive vibrational spectroscopic techniques have emerged as powerful alternatives to address these limitations (15, 17).

Molecular vibrations produce specific Raman spectra that can be used as a signature of molecules. Label-free Raman spectroscopy has been extensively applied to study cancer cell metabolism and classification (18). However, spontaneous Raman spectroscopy has limitations in biological imaging applications due to very low signal levels and slow image acquisition speeds. Coherent Raman Scattering processes, which include Coherent Anti-Stokes Raman Scattering (CARS) and Stimulated Raman Scattering (SRS), overcome these limitations and allow high-speed vibrational imaging of living specimens (19). CARS imaging has been used in studying cancer cell metabolism, but it suffers from a non-resonant background (15, 20). SRS imaging shows a better contrast as it does not have the problem of non-resonant background. SRS imaging has emerged as a proficient method for studying cell metabolism, including cancer cells, in the past decade. Applications of SRS imaging to cancer cell metabolism have revealed new cancer biomarkers (21, 22). Being able to generate maps of multiple biomolecules, recently developed high-content hyperspectral SRS (h^2^SRS) imaging shows great potential in profiling cancer cell metabolism in live and fixed samples (23, 24).

While various Raman techniques have been used to study cancer cells, CAR-T cells or other immune cells (25, 26), using coherent Raman microscopy to study the result of CAR-T cell challenge on cancer cell metabolism is under-explored. In this work, we studied the alteration of ovarian cancer cell metabolism using a CAR-T cell and cancer cell coculture. Towards this goal, we harnessed both near-infrared (NIR) and Visible h^2^SRS imaging (23, 24, 27, 28) to study lipid metabolic changes, especially the changes in cholesterol metabolism, in ovarian cancer cells cocultured with CAR-T cells. Based on the observation of increased cholesteryl ester storage and free cholesterol levels, we further explored the impact of cholesterol metabolic modulation on the efficiency of CAR-T cell-mediated killing of ovarian cancer cells.

## Results

### NIR h^2^SRS imaging shows altered lipid metabolism and protrusions in ovarian cancer cells co-cultured with CAR-T cells

To investigate the metabolic changes in the ovarian cancer cells under attack from CAR-T cells, we employed near-infrared (NIR) h^2^SRS imaging of a coculture of ovarian cancer and CAR-T cells. We cocultured an ovarian cancer cell line, SKOV3, with anti-HER2 CAR-T cells. We acquired NIR h^2^SRS imaging stacks of the coculture samples using a lab-built setup (27), shown in **Fig. 1A** and described in the methods section. As shown in **Fig. 1B**, we then performed spectral unmixing of these stacks using pixel-wise least absolute shrinkage and selection operator (LASSO) to decompose them into chemical maps of triacylglycerols/triglycerides, protein, total cholesterol, and nucleic acid based on the pure chemical spectra acquired of trioleate glycerol, bovine serum albumin (BSA), cholesterol, and purified ribonucleic acid (RNA) from cells, respectively (23, 27). **Fig. 1C** shows the metabolic maps of the SKOV3 cells with and without coculture with CAR-T cells at an Effector: Target (E:T) of 2:1. We observed an overall signal increase from the SKOV3 cells cocultured with CAR-T cells.

**Figure 1.**
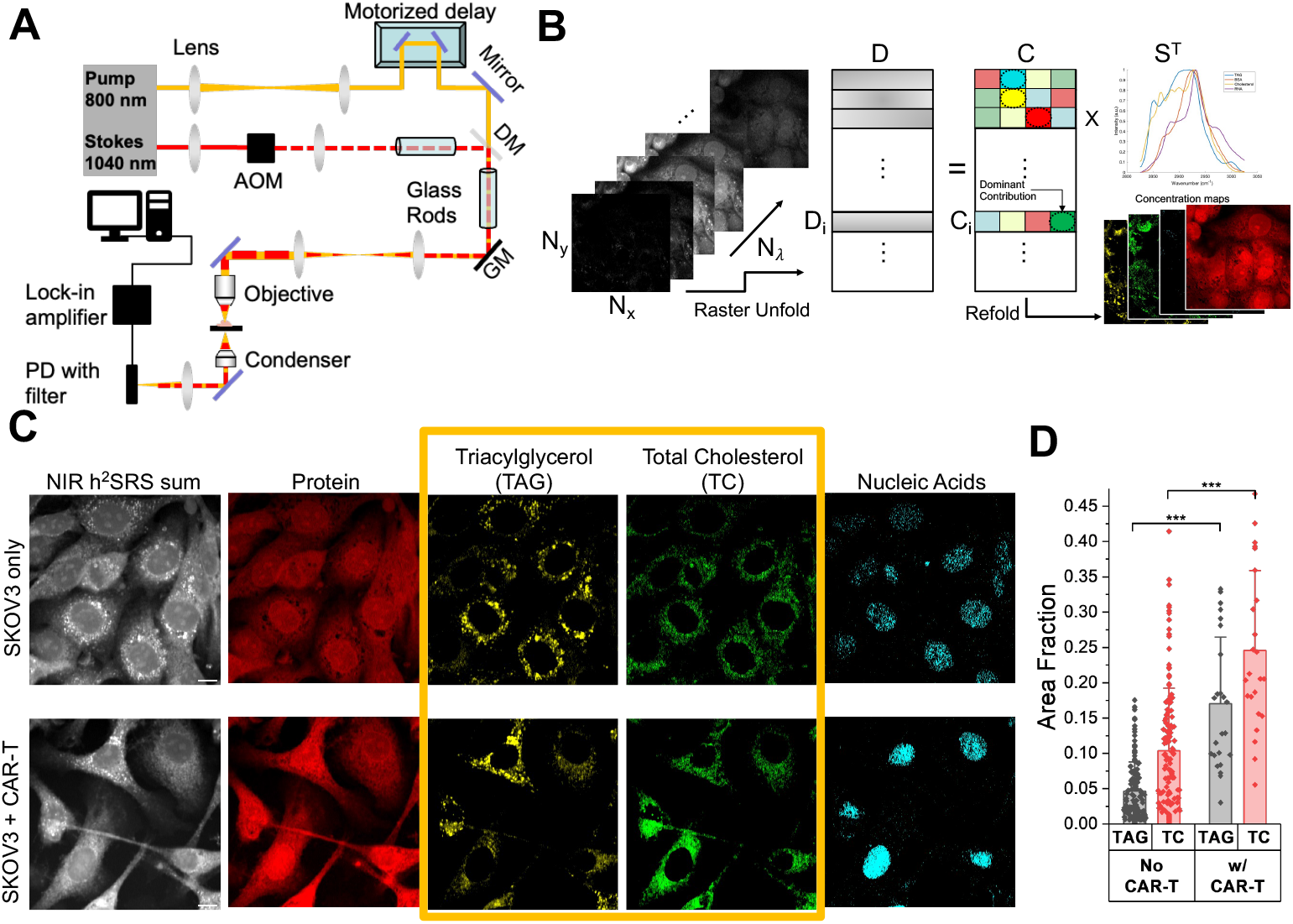
Altered lipid metabolism and protrusion in SKOV3 cells cocultured with CAR-T cells. **(A)** NIR SRS imaging setup. DM: Dichroic Mirror, GM: Galvo Mirror, AOM: Acousto-Optic Modulator, PD: Photodiode. **(B)** Schematic of spectral unmixing of hyperspectral images of dimensions, N_x_, N_y_, and N_y_ using pixel-wise least absolute shrinkage and selection operator (LASSO) into biochemical maps, D stands for data matrix, C for Concentration matrix and S for pure chemical spectra reference matrix. **(C)** NIR h^2^SRS imaging of the SKOV3 cells with and without CAR-T coculture, E:T = 2:1, scale bars: 10 μm. **(D)** Quantification of cellular Total Cholesterol (TC) and Triglycerides/Triacylglycerols (TAG) in SKOV3 cells with and without CAR-T coculture, n>30 per group, data presented as means +/+ stdev, *** p<0.001.

Furthermore, significant protrusions in the SKOV3 cells were observed in the coculture sample, indicating that the cells are under stress (29). Cell membrane protrusions are associated with cancer cell migration and metastasis (30, 31) and are also promoted in response to immune activity (32). Cancer cells under stress have been shown to increase the level of lipids and cholesterol species inside the cells as a survival mechanism (29, 33–37). Consequently, we analyzed the triglycerides (TAG) and total cholesterol (TC) maps shown in **Fig. 1C** to obtain quantitative results as presented in **Fig. 1D**. As expected, surviving SKOV3 cells cocultured with CAR-T cells exhibited an upregulation of triglycerides and total cholesterol (free cholesterol and cholesteryl ester), with increases of 3.5-fold and 2.3-fold, respectively, compared to SKOV3-only cells.

### Visible h^2^SRS imaging unveils upregulation of free cholesterol and cholesteryl ester in ovarian cancer cells cocultured with CAR-T cells

We further dissected the total cholesterol (TC) content to quantify free cholesterol (FC) and cholesteryl ester (CE), as these species are markers of distinct functions and regulations within the cell (35). Free cholesterol resides in membranes, while the core of lipid droplets (LDs) is mainly composed of TAG and sterol esters (38). High-resolution imaging to resolve single LDs and membranes would allow us to dissect the LDs content. Our recently developed Visible h^2^SRS system, shown in **Fig. 2A**, offers a diffraction-limited resolution of 86 nm (24, 39). Such Visible h^2^SRS imaging significantly improves organellar analysis compared to NIR h^2^SRS (24) since the size of lipid droplets can vary from less than 0.4 μm to tens of μm(38). As shown in **Fig. 2B**, Visible h^2^SRS resolved individual lipid droplets smaller than 0.2 μm, whereas in NIR h^2^SRS, the small LDs appear as aggregates. Thus, with Visible h^2^SRS, free cholesterol, cholesteryl ester, and TAG can be spatially resolved.

**Figure 2.**
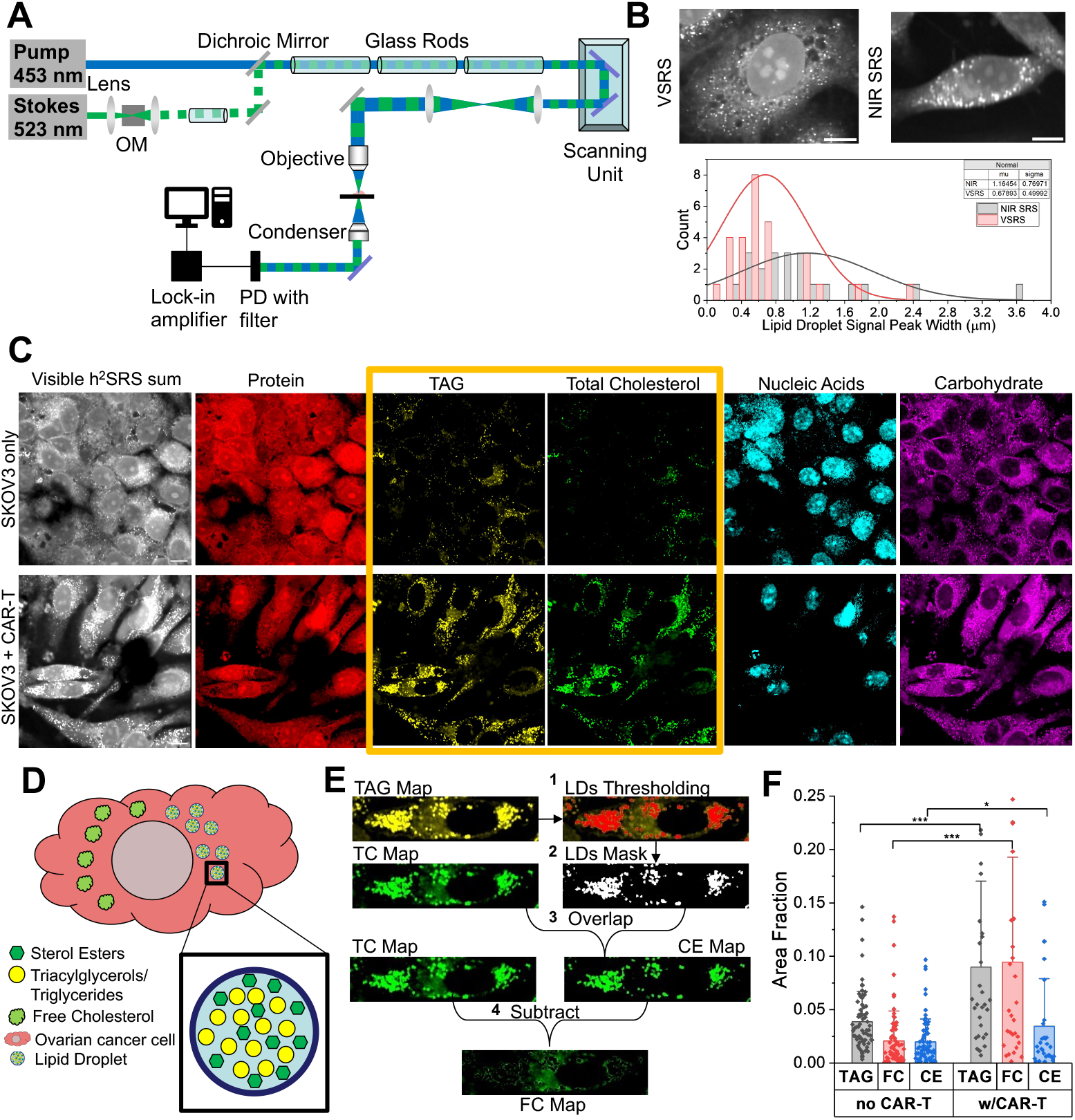
Visible SRS imaging uncovers altered cholesterol metabolism of SKOV3 cells. **(A)** Visible SRS imaging Setup, OM: Optical Modulator, PD: Photodiode. **(B)** Histogram of LDs spatial resolution comparison of NIR SRS and Visible SRS. **(C)** Visible h^2^SRS imaging of the SKOV3 cells with and without CAR-T coculture, E:T = 2:1, scale bars: 10 μm. **(D)** Composition of Lipid Droplets. **(E)** Schematic of CE and FC map generation from TAG and TC maps. **(F)** Quantification of TAG, FC and CE in SKOV3 cells with and without coculture with CAR-T cells., n>30 per group, data presented as means +/+ stdev, *** p<0.001, * p<0.05, TC: Total Cholesterol, FC: Free Cholesterol, CE: Cholesteryl Ester, TAG: Triacylglycerols/Triglycerides.

We acquired Visible h^2^SRS images of the SKOV3 cells only, and SKOV3 and CAR-T cells cocultured at an E:T ratio of 2:1. Metabolic maps are shown in **Fig. 2C**. We also acquired the reference spectra of triacylglycerol, protein, total cholesterol, carbohydrates, and nucleic acid using trioleate glyceride, bovine serum albumin (BSA), cholesterol, glucose, and purified ribonucleic acid (RNA) from cells, respectively for spectral unmixing. The symmetric CH_3_ stretching in both CE and cholesterol contributes to the Raman spectrum peak at 2870 cm^-1^ (40, 41). This peak contributes to the spectral unmixing in the same way, and it results in both CE and cholesterol being assigned to the same chemical map of total cholesterol. As previously mentioned, and shown in **Fig. 2D**, TAG and sterol esters make up most of the LDs. The cholesterol signal in the LD area should be from cholesteryl ester (38), which is the excess cellular cholesterol esterized for storage in LDs (33, 42). As shown in **Fig. 2E**, we can separate the CE map from the TC map by creating a droplet mask out of the TAG map and overlapping it with the TC map. The free cholesterol (FC) content was then calculated as the difference between TC and CE. We further analyzed the FC and CE area fractions in each cell of the two groups (SKOV3 only and SKOV3+CAR-T), as shown in **Fig. 2F**. We observed that all the lipid species were upregulated in the SKOV3 cells cocultured with CAR-T cells compared to the SKOV3-only cells, with specific increases of 2.3-fold in TAG, 4.5-fold in FC, and 1.7-fold in CE, respectively. Previous studies have shown that cholesterol plays a significant role in cancer and immune interactions (9, 12, 34, 43–45). The cholesterol level in the tumor microenvironment is usually elevated, contributing to cancer progression and affecting immune cell response (12, 43, 46, 47). Studies have shown that altering cholesterol levels in cancer cells can enhance T-cell function (44, 46). Both CE and FC increase in cancer cells have been linked to cancer cell aggressiveness, immune response blockade, survivability, and proliferation (44, 48, 49). Through chemical imaging of the surviving SKOV3 cells in the SKOV3+CAR-T coculture, we revealed an upregulation of intracellular CE and TC levels in response to CAR-T challenge.

### Inhibition of cholesterol synthesis or esterification suppresses SKOV3 mobility

To investigate the effects of lowering cellular levels of either CE or TC in SKOV3 cells, we employed Avasimibe and Simvastatin to intervene in cholesterol metabolism (50, 51). A previously published study showed that an increased level of free cholesterol in cancer cells softens cell membranes, facilitating cell migration (44) indicating that cholesterol metabolic intervention may reduce metastatic potential. As such, Simvastatin has been shown to decrease the cellular cholesterol level by inhibiting 3-hydroxy-3-methylglutaryl coenzyme A (HMG-CoA) reductase and, in turn, reducing ovarian cancer cell metastatic properties (13, 52). Simvastatin has also been studied for T cell response enhancement in solid cancers like lung, breast, and gastric cancer (53–55). Concurrently, CE accumulation is associated with the upregulation of the Phosphatidylinositol 3-Kinase/Protein Kinase B (PI3K/AKT) pathway connected to increased cancer aggressiveness and metastasis (48). The enzyme Acyl-CoA cholesterol acyltransferase 1 (ACAT-1) is responsible for cholesterol esterification of any excess cellular cholesterol (42, 48). Hence, inhibition of ACAT-1 by Avasimibe decreases cancer aggressiveness by inducing endoplasmic reticulum stress. Researchers have also demonstrated the role of ACAT-1 inhibition in T cells in enhancing T cell response against liquid (56, 57) and solid cancers (12).

As summarized in **Fig. 3A**, FC and CE are elevated in SKOV3 cells in the presence of CAR-T cells. While Simvastatin lowers the cellular FC, Avasimibe lowers the cellular CE. Hence, we examined the use of these drugs in decreasing the survival of SKOV3 cells cocultured with CAR-T cells. **Fig. 3B** shows the TAG and TC maps of the Visible h^2^SRS imaging of SKOV3 cells with and without treatment with Avasimibe and Simvastatin. The quantification of the images is presented in **Fig. 3C**. The SKOV3-only data presented in **Fig. 3C** are the same as in **Fig. 2F**, as these experiments were conducted in parallel. With Avasimibe treatment, the cellular CE level decreased to about 0.5-fold of the untreated group, while the FC and TAG levels stayed relatively the same. We also observed that with Simvastatin treatment, the cellular FC level decreased to about 0.25-fold of the untreated group, and the TAG levels increased 1.35-fold compared to the untreated group. However, as expected, cellular CE levels decreased (48) mildly (to 0.69-fold of the untreated group) following Simvastatin treatment, although the change was not statistically significant. With this validation, we conducted a migration assay with and without drug treatment of SKOV3 cells, presented in **Fig. 3D-G**. The concentrations used for the Avasimibe and Simvastatin treatments were at 30 μM and were selected based on the cell viability of greater than 80% using the dose response assay presented in Fig. S2C. We acquired confocal fluorescence images of the migrated cells in trans-well membranes. We can observe that the migration of the SKOV3 cells was significantly reduced following both Avasimibe and Simvastatin treatment by 95.5% and 47.4% compared to the untreated group. Importantly, Avasimibe treatment resulted in a much more significant reduction in migration, over twice that in the Simvastatin-treated group, showing that Avasimibe has greater potential for synergistic application with CAR-T therapy.

**Figure 3.**
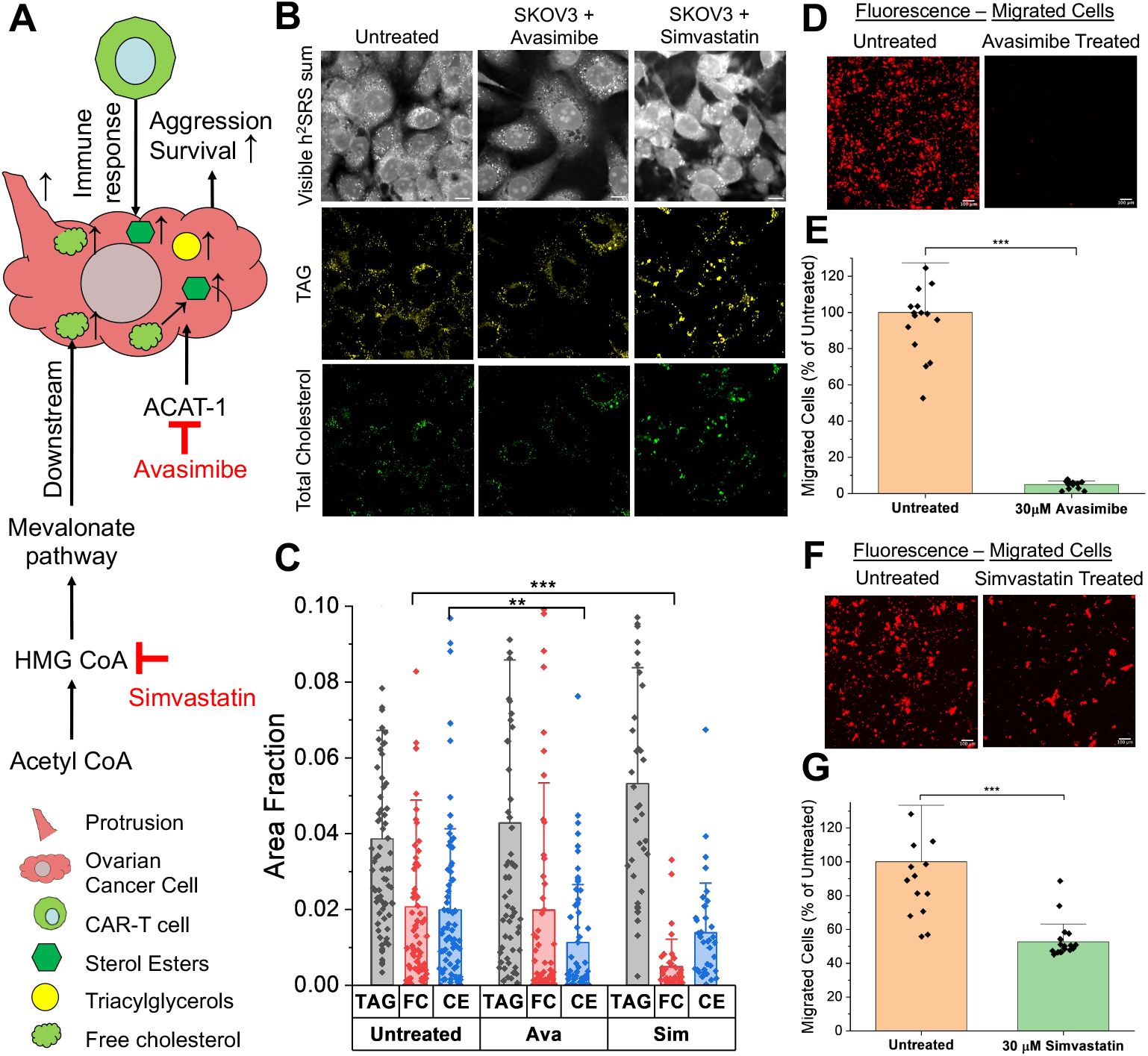
Drug intervention of cholesterol metabolism to reduce ovarian cancer aggression. **(A)** Summary of metabolism changes in ovarian cancer cells as a reaction to immune response and cholesterol metabolism intervention with Avasimibe and Simvastatin. **(B)** Visible h^2^SRS imaging TAG and TC maps of untreated, Avasimibe treated, Simvastatin treated SKOV3 cells. Avasimibe concentration: 30 μM, Simvastatin concentration: 10 μM, scale bars: 10 μm. **(C)** Quantification of cellular TAG, FC, CE in untreated, Avasimibe treated, Simvastatin treated SKOV3 cells, n>30 per group. **(D)** Confocal fluorescence imaging of migrated SKOV3 cells with Avasimibe and without Avasimibe. Avasimibe concentration: 30 μM, scale bars: 100μm. **(E)** Quantification of the migration with and without Avasimibe treatment, n=3 per group. **(F)** Confocal fluorescence imaging of migrated SKOV3 cells with Simvastatin and without Simvastatin, Simvastatin concentration: 30 μM, scale bars: 100μm. **(G)** Quantification of the cell migration with and without Simvastatin treatment, n=3 per group. Data presented as means +/+ std, *** p<0.001, ** p<0.01, TC: Total Cholesterol, FC: Free Cholesterol, CE: Cholesteryl Ester, TAG: Triacylglycerols/Triglycerides.

### Avasimibe treatment enhances CAR-T induced cancer cell killing in 2D cancer cell models

To evaluate the impact of combining metabolic treatment with CAR-T cell killing on ovarian cancer cells, we conducted cell killing assays on SKOV3 cells. We performed this assay on 96-well plates and read out measurements using a plate reader (Fig S4). For the two groups, we had untreated SKOV3 and treated SKOV3 cells. In the treated group, we incubated SKOV3 cells with 30 μM Simvastatin for 24 h. Then, both groups were followed by coculture with anti-HER2 CAR-T cells for 24 h at an E:T ratio of 1:2. The simvastatin treatment did not provide significant improvement in the CAR-T induced cytotoxicity in SKOV3 cells (**Fig. 4A**). Next, we carried out a cytotoxicity assay with Avasimibe, as presented in **Fig. 4B**. Based on prior evidence that Avasimibe can enhance T cell function (12, 56, 57), we included all combinations of Avasimibe treatment in this assay, treating either the cancer cells, the CAR-T cells, or both. The Avasimibe concentration used for SKOV3 treatment was 30μM for cell viability greater than 80%. For CAR-T cell treatment, the Avasimibe concentration was at 1μM (56). After 24 hours of Avasimibe treatment of SKOV3, the ovarian cancer cells were cocultured with untreated or Avasimibe-treated anti-HER2 CAR-T cells for another 24 hours at an E:T ratio of 1:4. The results showed that Avasimibe-treated SKOV3 cells, when cocultured with untreated CAR-T cells, exhibited higher cytotoxicity (54.6%) than the coculture group with untreated SKOV3 and untreated CAR-T cells (43.5%). Notably, treating only the SKOV3 cells led to greater cell death than treating both SKOV3 and CAR-T cells (51.6%) or only the CAR-T cells (46.8%). These results suggest that pretreatment of the ovarian cancer cells is sufficient to improve the CAR-T cell killing. We corroborated these results by adapting these ovarian cancer pre-treatment conditions to 35 mm glass bottom dishes and imaging using a lab-built widefield fluorescence setup described in the methods section. We incubated SKOV3 cells either with treatment of 30 μM Avasimibe or no treatment for 24 hours. We then added untreated anti-HER2 CAR-T cells at an E:T ratio of 2:1 for a 24-hour coculture. As shown in fluorescence images (**Fig. 4C**) and the corresponding quantification presented in **Fig. 4D**, the Avasimibe-treated SKOV3 group showed the highest CAR-T cell induced cytotoxicity (76.4%) than the untreated SKOV3 group (54.6%) each cocultured with untreated CAR-T cells.

**Figure 4.**
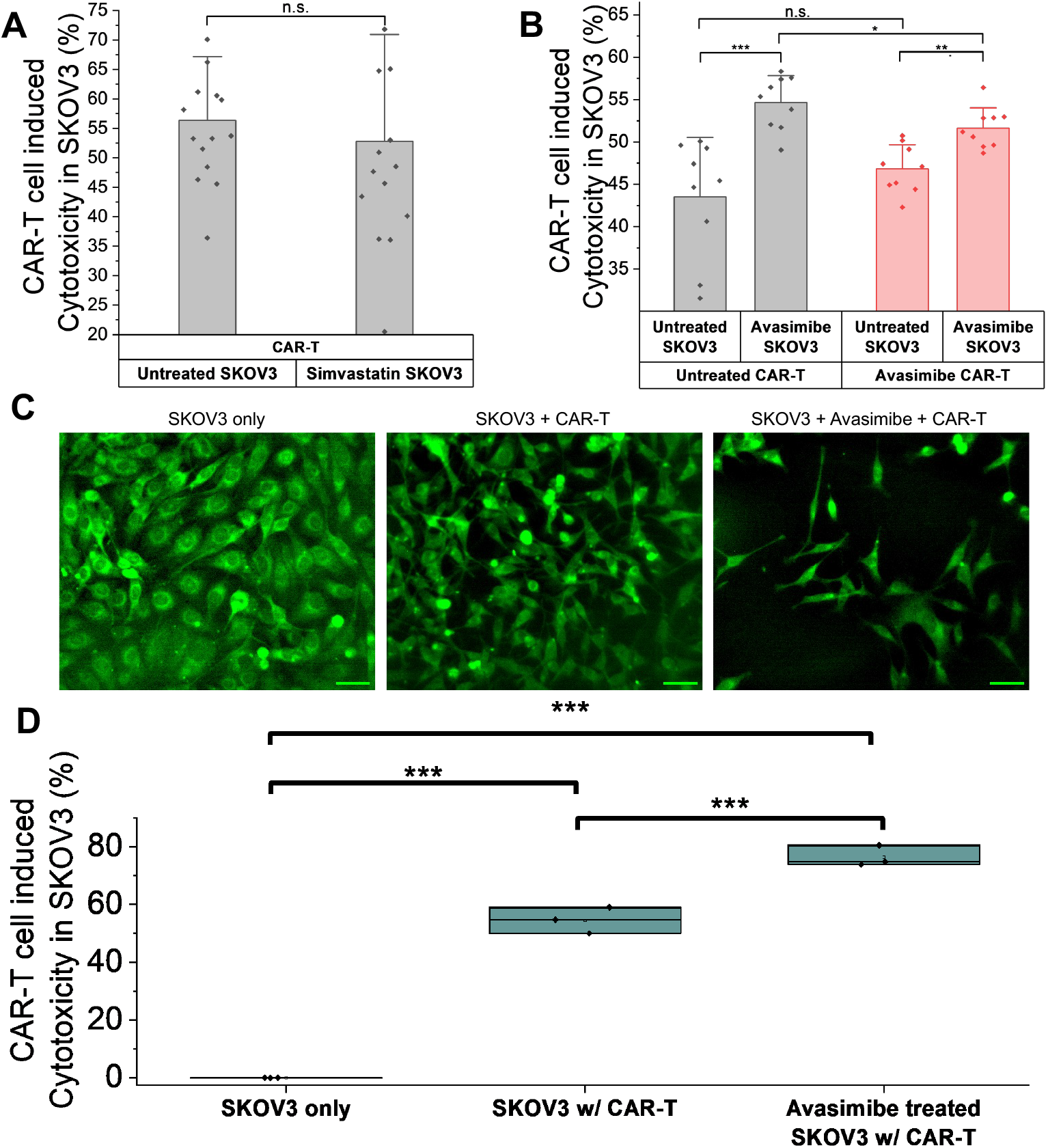
Drug intervention to enhance CAR-T cell induced cytotoxicity in ovarian cancer cells. **(A)** Cytotoxicity assay of untreated and Simvastatin treated SKOV3 cells with CAR-T cells and simvastatin treatment, n=15 per group, Simvastatin concentration: 30 μM. **(B)** Cell killing assay of untreated and Avasimibe treated SKOV3 cells with untreated and Avasimibe treated CAR-T cells, n=9 per group, Avasimibe concentration: 30 μM for SKOV3 cells and 1 μM for CAR-T cells. **(C)** Widefield fluorescence imaging of CAR-T-cell mediated cytotoxicity of the SKOV3 cells, scale bars: 100 μm. **(D)** Quantification of fluorescence imaging for the secondary cytotoxicity assay in C, n=3 per group. Data presented as means +/+ stdev, ***p<0.001, ** p<0.01, * p<0.05, n.s. non-significant, p>0.05.

### Avasimibe treatment enhances CAR-T induced cell killing in 3D tumor spheroids

We further extended the CAR-T cell induced cell killing assay to a 3D spheroids model. As has already been established in earlier studies, 3D spheroid models resemble the tumor micro-environment more closely than 2D cell cultures (58). We used GFP-SKOV3 cells to formulate 3D spheroids in ultra-low attachment 96-well plates. We incubated the GFP-SKOV3 cells in these plates for 48 h until the spheroids formed and then cultured them with or without Avasimibe for the next 24 h. We then added untreated anti-HER2 CAR-T cells in either a 1:1 E:T ratio as a reference condition or a 5:1 E:T ratio to assess enhanced cytotoxicity. We used the cytotoxicity index, the ratio of dead cells to the total dead and live cells in the spheroids, to analyze the effect on cell killing efficacy. The confocal fluorescence images of the spheroids shown in **Fig. 5A-E** clearly indicate a positive impact of Avasimibe treatment on CAR-T induced cytotoxicity. Furthermore, as quantified using a plate reader and shown in **Fig. 5F**, the cytotoxicity enhancement is the highest at 0.62 in the Avasimibe-treated spheroids cocultured with CAR-T cells in a 5:1 E:T ratio. The comparable cytotoxicity indices in the 1:1 E: T reference groups, 0.21 for Avasimibe-treated and 0.19 for untreated spheroids (with not a statistically significant difference), indicate that Avasimibe does not induce cytotoxicity in SKOV3 spheroids in the absence of robust CAR-T cell activity, at the same time, its potentiating effect is evident at higher E: T ratios. Taken together, these results indicate that Avasimibe pre-treatment of ovarian cancer cells improves the cytotoxicity induced by CAR-T cells.

**Figure 5:**
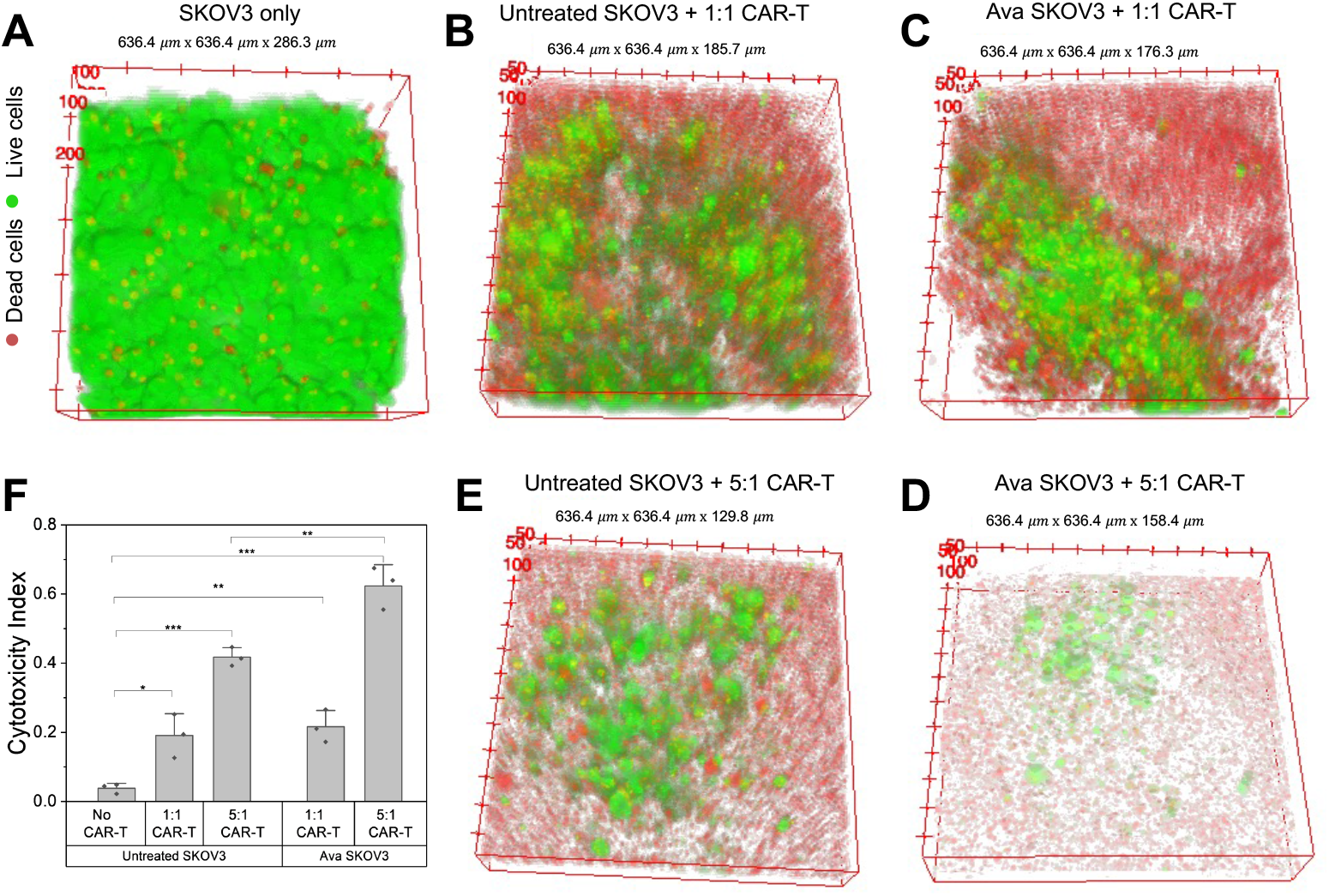
Avasimibe amplifies CAR-T induced cytotoxicity in 3D SKOV3 spheroids. **(A-E)** Confocal fluorescence imaging of GFP-SKOV3 cells spheroids incubated in various treatment conditions **(A)** Untreated, **(B)** Untreated spheroids incubated with CAR-T cells in E:T = 1:1, **(C)** Avasimibe treated spheroids incubated with CAR-T cells in E:T = 1:1, **(D)** Avasimibe treated spheroids incubated with CAR-T cells in E:T = 5:1, **(E)** Untreated spheroids incubated with CAR-T cells in E:T = 5:1. **(F)** Quantification of the spheroids cytotoxicity assay using plate reader, n=3 per group. Data presented as means +/+ stdev, ***p<0.001, ** p<0.01, * p<0.05.

### Cytokines impact lipid and cholesterol metabolism in cancer cells

The immune response by CAR-T cells involves a release of pro-inflammatory cytokines including Interleukin-2 (IL-2), Tumor Necrosis Factor-alpha (TNF-α), and Interferon-gamma (IFN-γ) (59). These cytokines have essential anti-tumor functions, including enhancing cytotoxic T-cell activity and inducing tumor apoptosis. On the other hand, it is also known that cytokines have myriads of functions that impact cancer cell growth and metabolism depending on the tumor microenvironment and immune state (60). To examine the molecular mechanism of cholesterol and triglycerides upregulation in ovarian cancer cells in the presence of CAR-T cells, we studied the impact of inflammatory cytokines on cancer cell metabolisms. As shown in **Fig. 6A**, we acquired NIR h^2^SRS images of SKOV3 cells incubated with complete culture medium with and without the supplementation of the inflammatory cytokines and focused on TAG and TC maps. We analyzed the TAG and TC maps (**Fig. 6B**), which revealed that both the fractions were upregulated in the SKOV3 cells treated with the cytokines, 2.3-fold and 2.94-fold with IL-2, 3.15-fold and 4-fold with TNF-α, 2.27-fold and 3.19-fold with IFN-γ, respectively. Notably, incubation with TNF-α showed the most impact among all the other cytokines. TNF-α has been shown to help in cancer progression and metastasis, which correlates with cholesterol upregulation in cancer cells (32, 61). With this knowledge, we conducted migration assays of SKOV3 cells cultured in culture medium with and without supplementation of the cytokines as shown in **Fig. 6C, D**. As expected, the TNF-α incubated cells showed the highest migration increase by nearly 100% compared to the untreated group. IFN-γ and IL-2 have complex roles in tumor progression and metastasis (59–61). In our results, IFN-γ treatment led to only a mild increase in migration (13.6%), while IL-2 treatment resulted in a more substantial increase (71.2%) relative to the untreated group. While pro-inflammatory cytokines such as TNF-α, IFN-γ, and IL-2 are known to have essential anti-tumor functions and are critical for immune activation, our results highlight that, under certain conditions, these same cytokines may paradoxically enhance cancer cell aggressiveness. Together, these findings suggest that cytokines might have context-dependent effects, potentially impacting the efficacy of CAR-T treatment.

**Figure 6.**
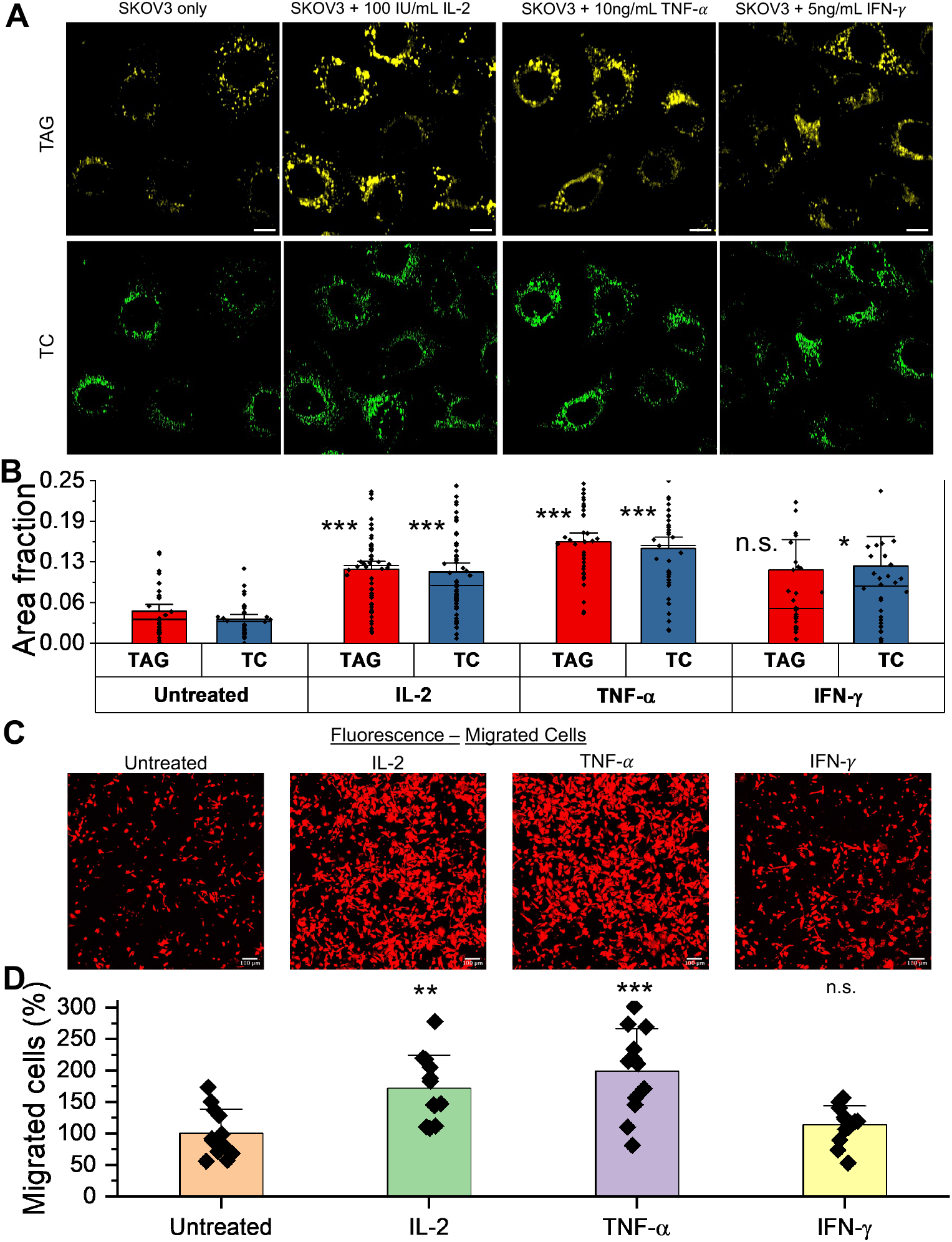
Impact of cytokines on metabolism and aggressiveness of SKOV3 cells. **(A)** TAG and TC NIR h^2^SRS imaging maps of SKOV3 cells with and without cytokines: IL-2, TNF-*α*, IFN-*γ* supplemented media for SKOV3 cells, IL-2 Concentration: 100 IU/mL, TNF-*α*Concentration: 10 ng/mL, IFN-*γ* Concentration: 5ng/mL, scale bars: 10 μm. **(B)** Quantification of cellular TC and TAG in SKOV3 cells incubated with medium with and without cytokines supplementation, n>31 per group. **(C)** Confocal fluorescence imaging of migrated SKOV3 cells with and without cytokines, scale bars: 100 μm. **(D)** Quantification of the cell migration with and without Cytokines treatment, n=3, *** p<0.001, ** p<0.01, data presented as means +/+ stdev, TC: total cholesterol. TAG: triglycerides/triacylglycerols.

## Discussion

CAR-T cell therapy is promising for liquid tumors, and its application in solid tumors, such as highly lethal ovarian cancer, is vital (1, 2). Numerous research studies have demonstrated distinct metabolic regulations within a tumor microenvironment (3, 8). Several studies have also shown that metabolic reprogramming is a crucial factor in cancer survivability and evasion of immune response (7, 10, 43). Despite these studies, there is a paucity of research on the direct metabolic impact of T-cell response on cancer cells. In this work, we employed NIR and visible h^2^SRS microscopy to investigate the metabolism modulation in ovarian cancer cells that survived the CAR-T cell attack. We noticed total cholesterol and triglyceride/triacylglycerol augmentation and protrusion development in the surviving ovarian cancer cells in the presence of CAR-T cells. These observable effects have been associated with increased cancer aggression and migration (13, 34, 42, 52), which might help cancer cells evade CAR-T cell response. Further, the dissection of lipid droplets using Visible h^2^SRS also showed an upregulation of the CE and FC within the cancer cells, suggesting a metabolic modulation as a survival mechanism (29, 33–37). This observation allowed us to choose drugs for metabolic intervention, helping enhance CAR-T cell anti-tumor response. Through various approaches, including a 3D spheroid culture model, we concluded that cholesteryl esterification inhibition using Avasimibe helps improve the CAR-T cell-induced cell killing in ovarian cancer.

CAR-T cells release pro-inflammatory cytokines, particularly IL-2, TNF-α, and IFN-γ, which can directly and indirectly affect tumor cells. These cytokines are critical for mediating anti-tumor immunity — IL-2 promotes the proliferation and persistence of cytotoxic T cells, IFN-γ enhances antigen presentation, and TNF-α can induce tumor cell apoptosis (59–62). However, these same cytokines can also contribute to tumor progression by promoting cancer cell survival, metabolic reprogramming, and migration, particularly in an inflammatory tumor microenvironment. Thus, we examined and validated that the lipid metabolic changes observed in surviving SKOV3 cells may result, at least partially, from cytokines among other reasons (29, 34–36). We further observed that cytokine supplementation increased ovarian cancer cell migration capacity. TNF-α led to the highest TC and TAG increase and a significant increase in ovarian cancer cell migration. This result is reasonable given that TNF-α, in addition to anti-tumor activity, is also known to have some pro-tumorigenic effects like cancer progression and migration by promoting epithelial to mesenchymal transition (32, 61), and the cancer migration is also linked to cholesterol upregulation in cancer cells (34). The cytokines IL-2 and IFN-γ also show pro-tumorigenic effects like immune exhaustion and cancer migration promotion (60, 62), although not as significant as TNF-α.

We observed that inhibiting cholesterol esterification in the cancer cells significantly improved the CAR-T cell-induced cytotoxicity against cancer cells compared to inhibition of cholesterol synthesis. CE accumulation is a sign of metabolic reprogramming in cancer, where excess cellular cholesterol and fatty acids are known to be stored in LDs as CE through cholesterol esterification catalyzed by ACAT-1 (33, 42) to avoid lipotoxicity and help with cancer cell survival under stress. Recent findings show that, more than the cholesterol component alone, fatty acids play a crucial role in cancer progression, membrane synthesis, and resistance to ferroptosis or oxidative stress (33, 42, 48, 63, 64). Cholesteryl esters often comprise arachidonic acid (AA), a polyunsaturated fatty acid, which is a key element in inflammation and is metabolized into compounds that have been shown to promote tumor progression, immune suppression, and resistance to therapy (48, 65). The LDs rich in CE, serving as lipid reservoirs, have shown to increase the metastasis and proliferation of the cancer cells. Diminishing the CE-rich lipid droplets by ACAT-1 inhibition using Avasimibe helps increase the cancer cell sensitivity through altered membrane dynamics and increased endoplasmic reticulum stress and reduces the cancer cell aggressiveness (42). Inhibiting the mevalonate pathway using statins has been shown to have a compensatory augmentation of SREBP activation and other ways of cholesterol upregulation (66). Thus, targeting cholesterol esterification may be more advantageous than targeting the cholesterol pathway directly.

Visible h^2^SRS uses visible lasers with extensive pulse chirping and noise suppression for imaging, which leads to a higher resolution than NIR h^2^SRS (24). This improvement in resolution allowed us to examine the lipid droplets. It was pivotal in analyzing lipid droplets, revealing changes in CE and FC levels in ovarian cancer cells, and offering insight into metabolic changes that support them. Thus, Visible h^2^SRS, with its high resolution and label-free chemical imaging inside the cells, holds much promise in studying diseases with metabolic dysregulations, especially in lipid droplets and cholesteryl esters. For example, in addition to other solid cancers, neurodegenerative diseases like Alzheimer’s, Parkinson’s, and Huntington’s diseases show altered lipid droplet dynamics and cholesterol metabolism (67). Exploring cellular metabolism in these diseases using Visible h^2^SRS would be beneficial to understand these dynamics, label-free, and for therapeutic discovery.

Although Visible h^2^SRS provides high spatial resolution, it has its limitations. We only studied the cell metabolism in the high-wavenumber (C-H) region. The C-H region compromises chemical specificity because of the presence and overlap of many signature peaks of chemicals in this region (27). High-content SRS imaging in the fingerprint region would provide a comprehensive observation of the metabolism changes in the pathways, not limited to only lipids (27, 38). Furthermore, Visible SRS imaging suffers from photodamage and limited penetration depth, which is unsuitable for biological tissue imaging. Imaging modalities, like NIR h^2^SRS in fingerprint region, provide more benefit in tissue imaging and facilitate access to more detailed metabolic information like amino acids, fumarate. Alternative methods using infra-red (IR) laser like, mid-infrared photothermal (MIP) microscopy also can be used to achieve submicron spatial resolution imaging and in the fingerprint region by detecting thermal expansion induced by IR using a visible probe (68), which has also been used to map the drug distribution successfully and might help in investigating how drug intervention helps enhance CAR-T cell induced cytotoxicity. These imaging modalities are promising tools that can be used to investigate metabolism modulation in various diseases, including but not limited to cancer.

## Materials and Methods

### Cell culture preparation

SKOV3 cells (purchased from American Type Culture Collection) for 2D culture and GFP-SKOV3 (generous gift from Dr. Matei Lab, Northwestern University) for 3D spheroids were cultured in the DMGM Basal Medium (Cell applications, MCBD 105 Medium and Corning, Medium 199 1X combined 1:1) supplemented with 10% fetal bovine serum (FBS) and penicillin/streptomycin (P/S; 100 U/ml). For the Anti HER2 CAR-T cells, normal whole peripheral blood from deidentified donors was obtained from Boston Children’s Hospital. Primary human CD3+ T cells were isolated from anonymous healthy donor blood by negative selection (STEMCELL, 15621). T cells were cultured in human T cell medium consisting of X-Vivo 15 medium (Lonza, 04418Q), 5% Human AB serum (Valley Biomedical, HP1022), 10 mM N-acetyl L-Cysteine (Sigma Aldrich, A9165), 55 μM 2-mercaptoethanol (Thermo Fisher Scientific, 31350010) supplemented with 50 units/mL IL-2 (NCI BRB Preclinical Repository). T cells were cryopreserved in 90% heat-inactivated FBS and 10% DMSO. All the cells were cultured and maintained in an incubator with 5% CO_2_ and 80-90% humidity at 37^0^C. We prepared coculture samples in an E:T ratio of 1:2 in 35 mm glass bottom dishes and fixed them with 10% formalin for h^2^SRS imaging. The coculture cell samples were prepared following the protocol shown in Fig. S1. For the cytokines treated imaging, we cultured SKOV3 cells with complete culture medium supplemented with 100 IU/mL IL-2 (NCI BRB Preclinical Repository), 10 ng/mL TNF-*α* (Thermo Fisher Scientific, PHC3011), 5 ng/mL IFN-*γ* (Thermo Fisher Scientific, PHC4033) respectively for 24 h. The cells were then fixed with a 10% formalin solution before imaging.

### Lentiviral Transduction of Human T Cells

Second-generation lentivirus was packaged via transfection of lentixHEK 293 cells (Takara) with a pHR transgene expression vector and the viral packaging plasmids: pMD2.G encoding for VSV-G pseudotyping coat protein (Addgene, 12259), pDelta 8.74 (Addgene, 22036), and pAdv (Promega). One day after transfection, viral supernatant was harvested every 24 hours for 3 days and replenished with pre-warmed FreeStyle 293 expression media (Gibco) with 2 mM L-glutamine, 100 U/ml penicillin, 100 ug/mL streptomycin, 1 mM sodium pyruvate, and 5mM sodium butyrate. Then, 4X PEG8000 lentivirus concentrator was added to the harvested virus, and the mixture was incubated overnight before spinning at 1600xG for 50 minutes. Primary T cells were thawed 2 days before virus purification and cultured in the T cell medium described above. One day before virus centrifugation, T cells were stimulated with Human T-activator CD3/CD28 Dynabeads (Thermo Scientific #11132D) at a 1:1 cell-to-bead ratio and cultured for 24 hours. After viral supernatant purification, rectronectin (Clontech #T100B) was used to mediate T-cell transduction. Briefly, non-TC-treated 6-well plates were coated with rectronectin following the supplier’s protocol. Then, concentrated viral supernatant was added to each well and spun for 90 min at 1200xG. After centrifugation, the viral supernatant was removed, and 4 mL of human T cells at 250k/ml in T cell growth media supplemented with 100U/ml of IL-2 was added to the well. Cells were spun at 1200xG for 60 minutes and moved to an incubator at 37°C.

### Cell viability assay

The cell viability assay was conducted using the MTS reagent (Abcam, ab197010) in 96 well plates. SKOV3 cells were seeded in 96 well plates at the density of 5,000-15,000 cells per well and incubated for 24 hours, followed by the addition of the drug at the mentioned concentrations and incubated for another 24 hours. Then, the MTS reagent was added to the plate so that the culture medium to MTS reagent ratio was 5:1, and the mixture was incubated for 1h. The absorbance was then read at 490 nm using a plate reader (i3X SpectraMax Molecular Devices). The step-by-step protocol is shown in Fig. S2A. The treatment concentrations were selected so that they were sublethal (>80% viability) to study cell responses without causing immediate cell death, as shown in Fig. S2 (B, C). This criterion also applies to cytokines as they did not show significant toxicity to the SKOV3 cells, as shown in Fig. S2B.

### Migration assay and confocal fluorescence imaging

Migration assays were conducted using trans-well membranes (Corning, CLS3401). SKOV3 cells were seeded onto 35 mm cell culture dishes till 70% confluence. Then, the drugs at the mentioned concentration were added to respective dishes, and the cells were incubated for 24 hours. A culture medium supplemented with 20% FBS was added to the wells in 24 well plates, and the drugs at the mentioned concentrations. The cells were then detached from the dishes and seeded onto the upper chamber of the trans well membranes with culture medium only and no FBS. The drugs at respective concentrations were added to the top chamber. The cells were then incubated for 24 hours, followed by fixation with 10% formalin. The upper chambers of trans-well membranes were then cleaned with cotton swabs to remove non-migrated cells. The migrated cells were then dyed with propidium iodide for 30 minutes at a concentration of 50*μgmL*^−1^. We acquired confocal fluorescence images (excitation 561 nm, emission 570-620 nm) using a laser scanning confocal microscope (Olympus FV3000, Micro/Nano Imaging Facility, Boston University) and quantified them using ImageJ. Drugs used were Avasimibe, purchased from Selleck Chemicals, and Simvastatin, purchased from Thermo Fisher Scientific. A step-by-step protocol is shown in Fig. S3.

### CAR-T cell cytotoxicity assays

48 hours before assay, SKOV3 cells were seeded in 96-well plates at a density of 15,000-20,000 cells per well and incubated for 24 hours. Then, drugs were added at the mentioned concentrations, and the cells were incubated with dosed media for another 24 hours. 24 hours before assay, anti- HER2 CAR-T cells in their respective culture media were treated with drugs at the mentioned concentrations, if any. On the day of assay, SKOV3 cells were incubated with the green cell proliferation fluorescent agent (Abcam, ab176735) following the manufacturer’s protocol before coming into contact with CAR-T cells. Anti-HER2 CAR-T cells in various drug treatment conditions were added to SKOV3 cells at an appropriate E:T ratio. The coculture was incubated overnight before fluorescence read (excitation 488 nm, emission 511-525 nm), using a plate reader to assess target cell viability. A step-by-step protocol is shown in Fig. S4. A minimum of 9 wells per group were analyzed. For the secondary imaging validation, SKOV3 cells were seeded in the glass-bottom dishes at a density of 25,000 to 50,000 cells per dish for 24 hours. Then, the cells were incubated with Avasimibe at the specified concentration in the culture medium for 24 hours. Before the assay, the cells were incubated with the green proliferation dye following the manufacturer’s protocol. Anti-HER2 CAR-T cells in their respective culture medium were added at the E:T ratio of 2:1 and incubated for 24 hours. At the end of the coculture assay, the cells were fixed with 10% formalin before acquiring 6 fluorescence images per dish with our lab-built microscope using an Olympus IX71 microscope frame and 20x air objective (excitation 470 nm, emission 500-540 nm) (69). The images were analyzed using ImageJ.

For the 3D spheroids cytotoxicity assay, GFP-SKOV3 cells were seeded in ultra-low attachment round-bottom 96-well plates at a density of 8,000 cells per well and incubated for 48 hours till the spheroids formed. The drugs were then added at the mentioned concentrations, and spheroids were incubated with dosed media for another 24 hours. Anti-HER2 CAR-T cells in their respective culture media were then added to the CAR-T treatment spheroids group in the appropriate ratios and cultured for 24 hours. The dead cells in the spheroids were then dyed with propidium iodide for 1 hour at a concentration of 5 *μgmL*^−1^. We acquired confocal fluorescence images (excitation 561 nm, emission 570-620 nm for PI; excitation 488 nm, emission 500-540 nm for GFP) using a laser scanning confocal microscope (Olympus FV3000, Micro/Nano Imaging Facility, Boston University) and acquired fluorescence reads (excitation 535 nm, emission 617 nm for PI; excitation 488 nm, emission 510 nm for GFP) using a plate reader (i3X SpectraMax Molecular Devices) for the quantification. A step-by-step protocol is shown in Fig. S5.

### High-content stimulated Raman scattering imaging

High Content SRS (h^2^SRS) imaging is a method involving hyperspectral Stimulated Raman Scattering (hSRS) imaging followed by denoising and spectral unmixing of the hyperspectral data using pixel-wise least absolute shrinkage and selection operator (LASSO) into biochemical maps (23, 24, 27, 28). In this project, we used Near-Infrared (NIR) hSRS and Visible hSRS.

### NIR hSRS imaging setup

Our lab-built NIR hSRS system (27) is based on a dual-output laser (Insight DeepSee+, Spectral Physics), as shown in **Fig. 1A**. Hyperspectral imaging was achieved using the spectral focusing method by the temporal delay between pump and stokes beams, where both the femtosecond beams were chirped using glass rods to generate picosecond pulses and take hyperspectral stack image (27). We acquired an hSRS image stack for the cells in the CH wavenumber region in the range from 2820 cm^-1^ to 3050 cm^-1^ with Stokes beam (wavelength, 1040 nm) at 150mW and Pump beam (wavelength, 800 nm) at 30 mW measured before the galvo mirror.

### Visible hSRS imaging setup

Our lab-built visible hSRS system (24) is based on two broadband femtosecond visible lasers, as shown in **Fig. 2A**. The setup utilized a dual-output laser (Insight X3, Spectra Physics) where the output wavelengths of 906 nm and 1045 nm were used to obtain 453 nm and 523 nm through second harmonic generation. Hyperspectral imaging was achieved using the spectral focusing method, as mentioned previously in the NIR hSRS setup and detailed in (24). We acquired the hyperspectral imaging data for the cells in the CH wavenumber region as described above with Stokes beam (wavelength, 523 nm) at 30 mW and Pump beam (wavelength, 453 nm) at 20 mW measured before the galvo mirror.

### Self-Supervised Elimination of Non-Independent Noise (SPEND) denoising

We applied Self-Supervised Elimination of Non-Independent Noise (SPEND) denoising to the Visible SRS hyperspectral data to improve the resolution further. SPEND is a self-supervised deep learning denoising framework for removing non-independent noises in hyperspectral images. Details can be found in (70). In this study, the SPEND model was trained on an NVIDIA RTX 4090 GPU with 24 GB of memory, requiring approximately 1 hour for competition. The input dataset contains 24 fields of view. Each field of view contains 400 X 400 X 100 pixels. Data augmentation was performed using flipping and rotation, resulting in a fourfold increase in the effective dataset size. During interference, the model processes each field of view in 2.4 seconds.

### LASSO spectral unmixing and map analysis

Then, we used the pixel-wise LASSO method for the spectral unmixing of the hyperspectral stack into chemical maps of lipid, protein, cholesterol, and nucleic acid (27). It is a least square fitting problem with L1-norm regularization with the parameter *λ* used to control each chemical component’s sparsity level. The regularization is based on the observation that at each pixel, only a few chemical components contribute dominantly (27). The unmixing is implemented using acquired hSRS spectra for pure solutions of protein (Bovine Serum Albumin, BSA), lipid (Trioleate Glycerol, TAG), cholesterol, and nucleic acid (RNA solution) and carbohydrates (Glucose) as reference inputs for the operator wherever specified (Spectra shown in Fig. S6). The LASSO parameter, *λ*, was as follows for NIR h^2^SRS: *λ*_*TAG*_ = 0.05, *λ*_*protein*_ = 0.05, *λ*_*cholesterol*_ = 0.09, *λ*_*RNA*_ = 0.09. The LASSO parameter, *λ*, was as follows for Visible h^2^SRS: *λ*_*TAG*_ = 0.02, *λ*_*protein*_ = 0.03, *λ*_*cholesterol*_ = 0.0475, *λ*_*RNA*_ = 0.01, *λ*_*Glucose*_ = 0.02 or a single = 0.01 for all biochemical maps. For the total cholesterol (TC) and triglycerides/triacylglycerol (TAG) map analysis generated using both NIR and Visible h^2^SRS, we used Fiji (ImageJ). We quantified the TC and TAG in NIR h^2^SRS maps using ImageJ by applying a threshold based on intensity and calculating the area fraction for each out of the total cell area (33). We used the same method for the cholesteryl ester (CE), free cholesterol (FC) and TAG quantification using Visible h^2^SRS maps. To generate cholesteryl ester maps, we first generated a mask using TAG maps by applying a threshold to pick up only the LDs. Then, we calculated the overlap between the TAG masks and TC maps to get a cholesteryl ester map. Then, we calculated the FC fraction as a difference between TC and CE. We analyzed a minimum of 30 cells per group for statistical analysis.

## Statistical analysis

The statistical graphs were shown as means +/+ stdev unless specified otherwise. The statistical analysis was done using either unpaired t-test between treated and untreated groups and using OriginPro. Statistical significance was denoted as * for p<0.05, ** for p<0.01, *** for p<0.001, and n.s. for p>0.05 indicating a non-significant statistical difference.

## Acknowledgments

This work was funded by NIH grants R35GM136223 and R01CA224275 to J.X.C. Research reported in this publication was supported by the Boston University Micro and Nano Imaging Facility and the Office Of The Director, National Institutes of Health of the National Institutes of Health under Award Number S10OD024993. The content is solely the responsibility of the authors and does not necessarily represent the official views of the National Institute of Health. We thank Dr. Daniela Matei, Dr. Ana Maria Isaac, and Dr. Hao Huang from the Department of Obstetrics and Gynecology, Feinberg School of Medicine, Northwestern University (Chicago, IL 60611), for generously providing the GFP-SKOV3 cells. Their support was invaluable to this study.

## Author contributions

C.V.P.D., J.X.C and W.W. codesigned the experiments. C.V.P.D., J.H., H.L. performed the experiments, C.V.P.D and J.H. did the data analysis, M.Z., M.S., G.D. and H.H. helped in the data analysis. C.V.P.D. wrote the manuscript. All authors read and edited the manuscript.

## Notes

### Competing Interest Statement

The authors have declared no competing interest.

