## Supplementary figures for "Cholesterol metabolism modulation facilitates CAR-T induced killing of ovarian cancer"

### Supplementary Information

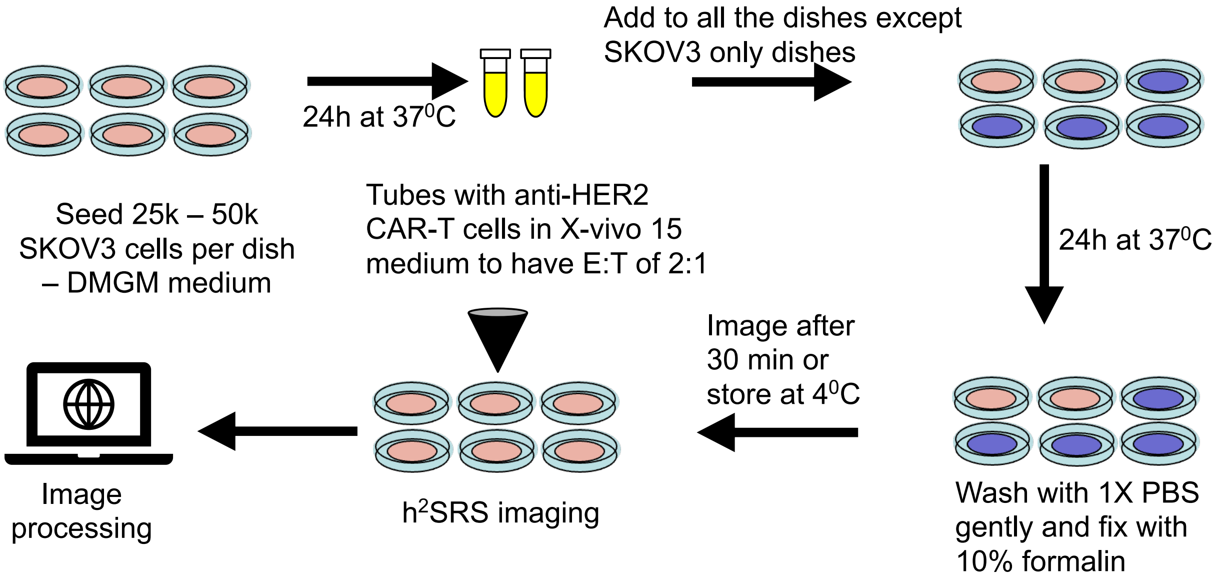


**Fig. S1. Coculture.** Coculture sample preparation for hSRS.

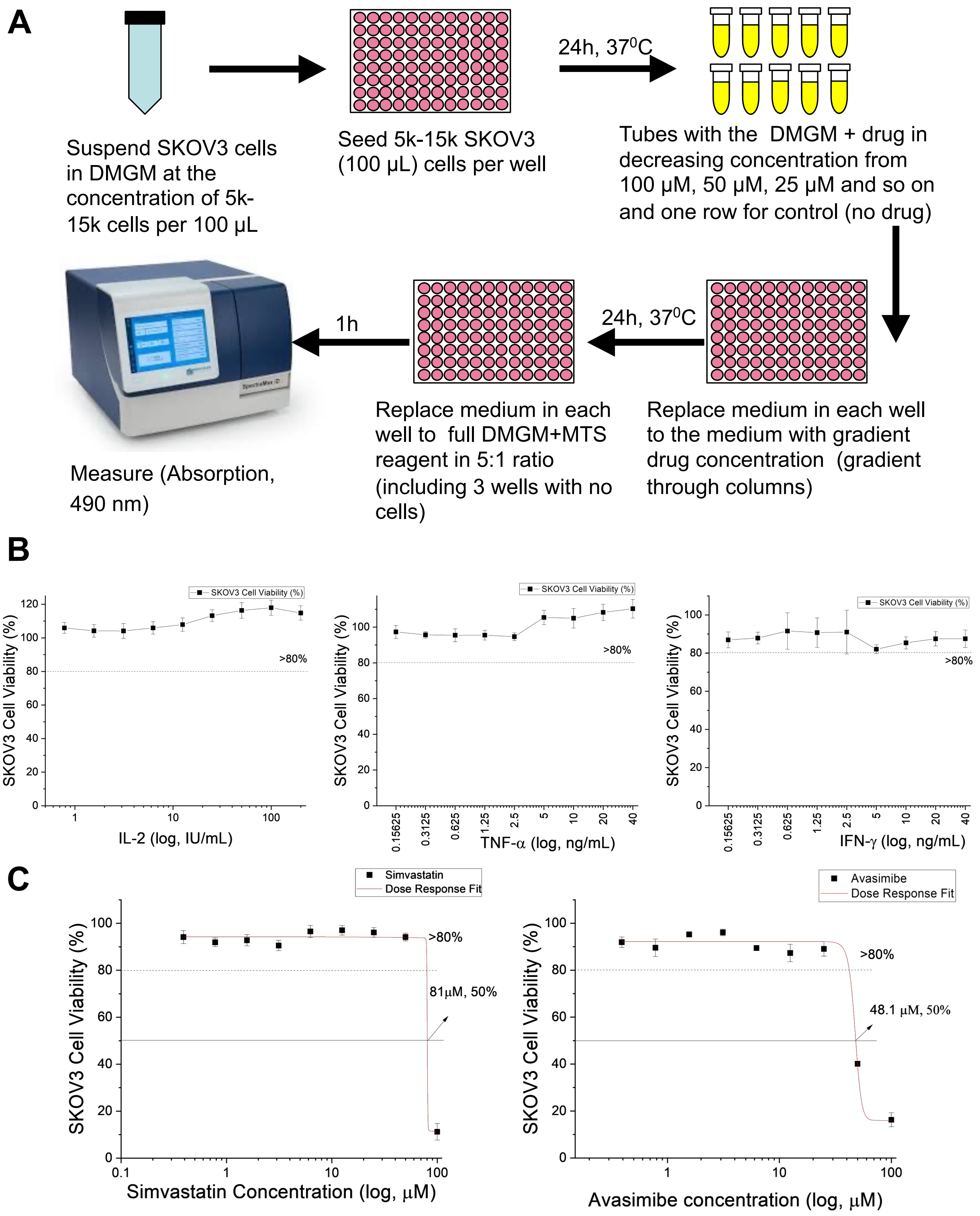


**Fig. S2. Cell viability assay protocol and dose response curves. (A)** Cell Viability Assay protocol with MTS Reagent. **(B)** Cell dose response for pro-inflammatory cytokines: IL-2, TNF-$\alpha$, IFN-$\gamma$ on SKOV3 cells, n=6 for each curve. **(C)** Cell dose response for Avasimibe and Simvastatin on SKOV3 cells, n=6 for each curve.

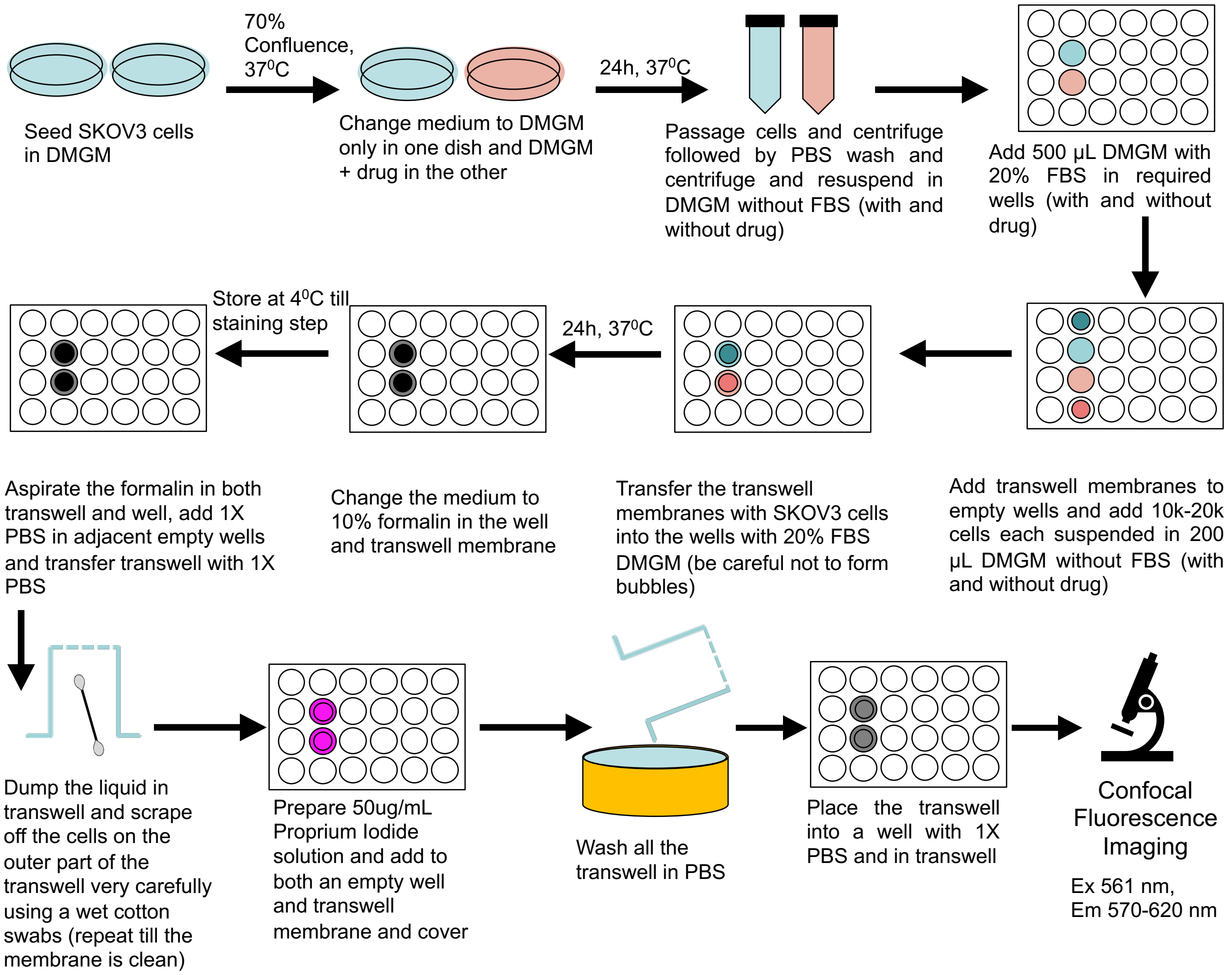


**Fig. S3. Migration Assay.** Migration assay protocol with and without drug treatment.

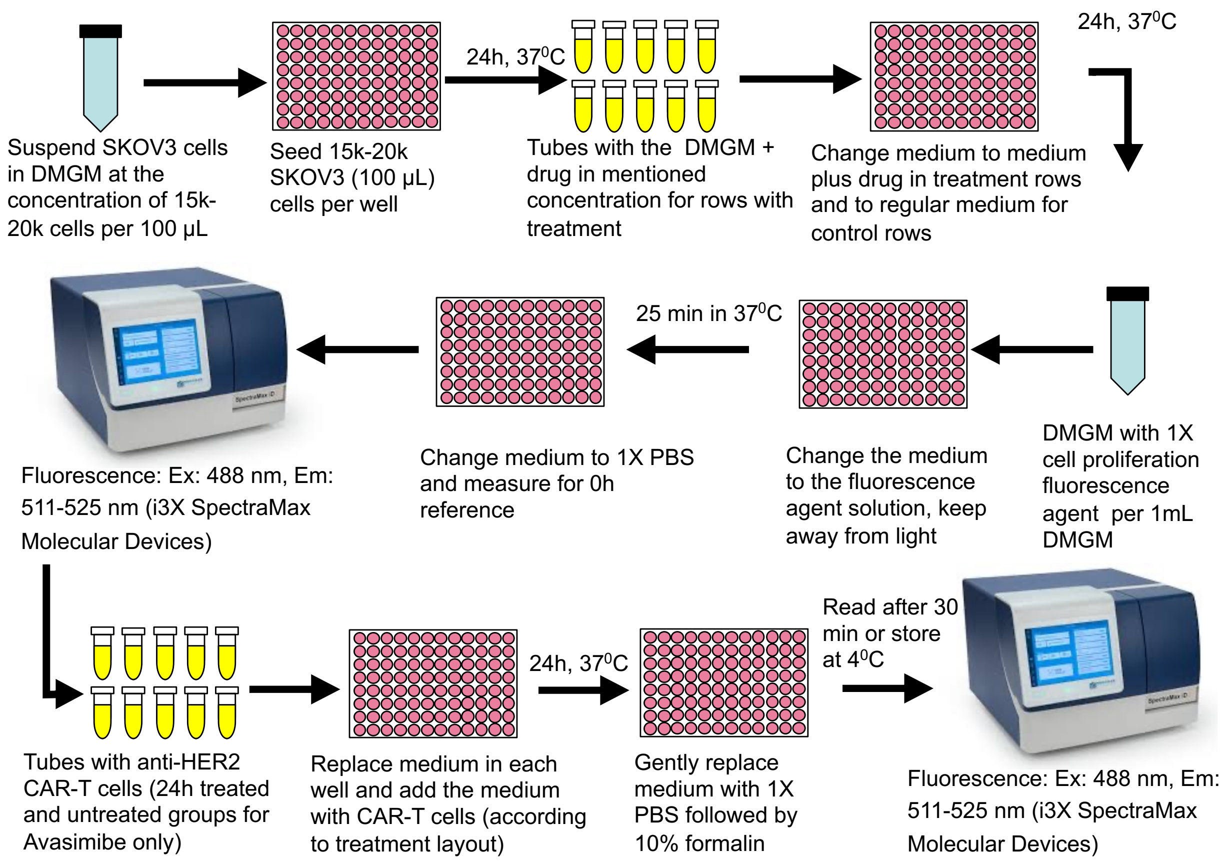


**Fig. S4. Cytotoxicity Assay.** CAR-T cell induced cytotoxicity assay protocol in 2D cultures.

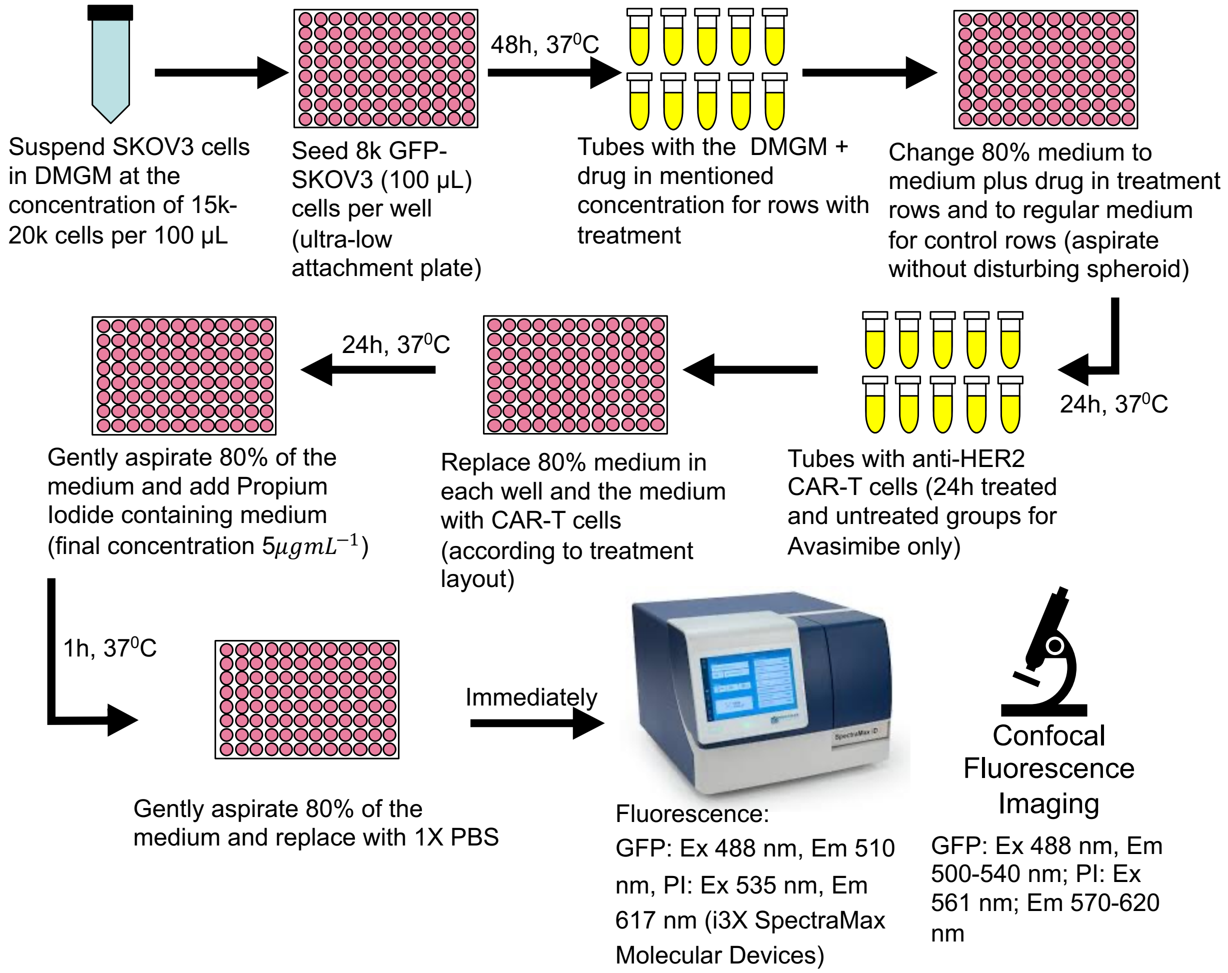


**Fig. S5. 3D spheroids cytotoxicity Assay.** CAR-T cell induced cytotoxicity assay protocol in 3D cancer spheroids.

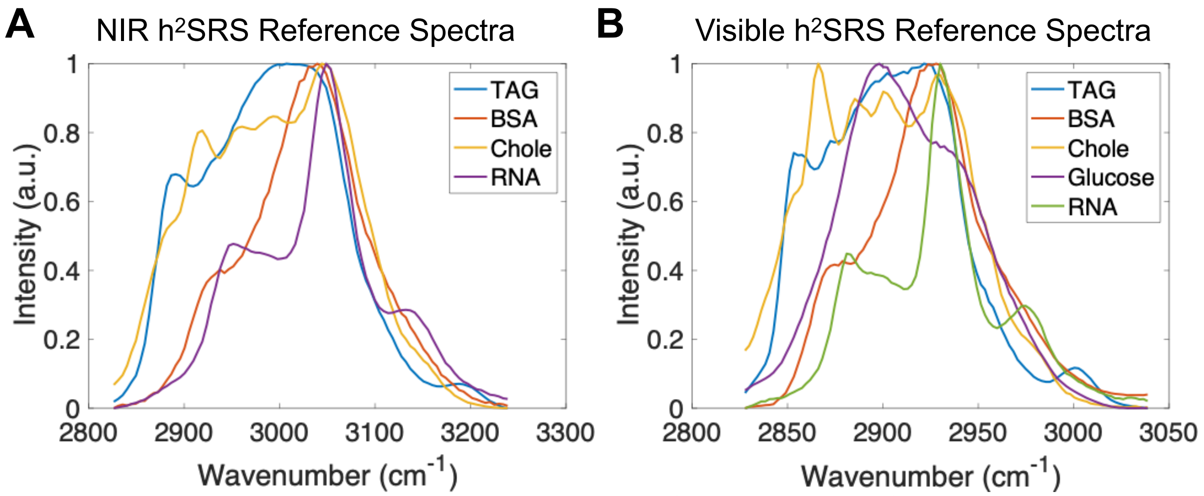


**Fig. S6. Pure Chemical Spectra Used for input in h$^2$SRS. (A)** NIR h$^2$SRS Reference Spectra, **(B)** Visible h$^2$SRS Reference Spectra
